# SOCS3 limits endotoxin-induced endothelial dysfunction by blocking a required autocrine interleukin 6 signal in human umbilical vein endothelial cells

**DOI:** 10.1101/2022.04.25.489373

**Authors:** Nina Martino, Ramon Bossardi Ramos, Dareen Chuy, Lindsay Tomaszek, Alejandro P Adam

## Abstract

Increased circulating levels of soluble interleukin (IL)-6 receptor α (sIL-6Rα) are commonly observed during inflammatory responses, allowing for IL-6 signaling to occur in cells that express the ubiquitous receptor subunit gp130 but not IL-6Rα, such as endothelial cells. Activation of Toll-like receptor (TLR)-4 or the tumor necrosis factor (TNF) receptor leads to NF-κB-dependent increases in endothelial IL-6 expression. Thus, we hypothesize that danger signals may induce autocrine IL-6 signaling within the endothelium via sIL-6Rα-mediated trans-signaling. In support of this hypothesis, we recently demonstrated that conditional deletion in the endothelium of the IL-6 signaling inhibitor SOCS3 leads to rapid mortality in mice challenged with the TLR-4 agonist endotoxin through increases in vascular leakage, thrombosis, leukocyte adhesion, and a type I-like interferon response. Here, we sought to directly test a role for sIL-6Rα in LPS-treated human umbilical vein endothelial cells. We show that cotreatment with sIL-6Rα dramatically increases the loss of barrier function and the expression of COX2 and tissue factor mRNA levels induced by LPS. This co-treatment led to a strong activation of STAT1 and STAT3 while not affecting LPS-induced activation of p38 and NF-κB signaling. Similar results were obtained when sIL-6Rα was added to a TNF challenge. JAK inhibition by pretreatment with ruxolitinib or by SOCS3 overexpression blunted LPS and sIL-6R synergistic effects, while SOCS3 knockdown further increased the response. Together, these findings demonstrate that IL-6 signaling downstream of NF-kB activation leads to a strong endothelial activation and may explain the acute endotheliopathy observed during critical illness.

## Introduction

Severe systemic inflammatory reactions often lead to multiorgan dysfunction, in part through an acute release of cytokines -a process called ‘cytokine storm’ (1). These cytokines and other inflammatory mediators promote vascular dysfunction leading to shock, including vasodilation, thrombosis, leukocyte plugging and systemic vascular leak. These defects result in severe hypotension, organ hypoperfusion and ultimately organ failure and death (2–4). IL-6 binding to a heterodimeric receptor that comprises IL-6Rα and gp130 activates the Janus tyrosine kinases (JAK) to initiate several downstream signal transduction cascades, including the activation of STAT3-dependent gene expression through its phosphorylation on tyrosine 705 (5, 6). Multiple positive and negative feedback loops regulate the intensity and duration of this signal. Chiefly, STAT3 promotes IL-6 and its own expression, thereby amplifying the signal. Concurrently, STAT3 mediates SOCS3 expression, leading to a potent negative regulation through SOCS3 binding to the IL-6R/JAK complex to block STAT3 activation (7).

While the endothelium does not express IL-6Rα (8, 9), the circulating levels of a soluble form (sIL-6Rα) quickly increase during acute inflammation, allowing the binding of IL-6 and IL-6Rα to gp130 molecules expressed by the endothelium to promote downstream signaling in a process termed ‘trans-signaling’ (5, 10). This mode of activation of IL-6 signaling in cultured human umbilical vein endothelial cells (HUVEC) is sufficient to promote a sustained barrier function loss (9). It is well accepted that activation of Toll-like receptor (TLR)-4 or the tumor necrosis factor (TNF) receptor leads to NF-κB-dependent increases in IL-6 expression (11), raising the possibility that danger signals may induce an autocrine IL-6 signaling within the endothelium. We hypothesize that circulating sIL-6Rα amplifies the endothelial response during acute inflammation initiated by innate immunity activation. In support of this hypothesis, we recently demonstrated (12) that conditional deletion in the endothelium of the IL-6 signaling inhibitor SOCS3 leads to rapid mortality in mice challenged with the TLR-4 agonist endotoxin (lipopolysaccharides from *E Coli*, LPS) through increases in vascular leak, thrombosis and leukocyte adhesion. We hypothesized that, in response to LPS, loss of SOCS3 specifically in the endothelium led to a strong autocrine signaling within the endothelium mediated by IL-6. Autocrine signaling involving interferons have been previously described in response to TNF signaling (13, 14).

To gain further mechanistic insight, we sought to directly test a role for IL-6 receptor in LPS-treated HUVEC. We show that a co-treatment with sIL-6Rα dramatically promoted endothelial dysfunction markers in response to LPS, including increases in COX2 and tissue factor mRNA levels and a large loss of monolayer barrier function. This co-treatment led to a strong activation of STAT1 and STAT3 while not affecting LPS-induced activation of p38 and NF-κB signaling. Similar results were obtained when sIL-6Rα was added to a TNF challenge. Consistent with our observations in endotoxemic mice lacking endothelial SOCS3 expression, this response in HUVEC was increased by SOCS3 knockdown and blunted by SOCS3 overexpression. Together, these findings demonstrate that IL-6 signaling downstream of NF-kB activation leads to a strong endothelial activation and may explain the acute endotheliopathy observed during critical illness. Moreover, blocking the IL-6 response in endothelial cells may allow for novel therapeutic strategies to blunt endothelial dysfunction during acute inflammatory responses.

## Methods

### Reagents and antibodies

Vendor, catalog numbers and concentrations used for reagents are listed in the supplemental table 1. Antibodies, including vendor catalog numbers, RRIDs and the specific concentrations and blocking solutions used are listed in the supplemental table 2.

### Cell culture and treatments

HUVECs were isolated in-house according to established protocols (9, 15–17) from anonymized healthy donors following the recommendations of the Albany Medical Center IRB. There was no exclusion of donors based on race, newborn sex, or age of mother. Umbilical cords (20–30 cm) from scheduled cesarean sections were stored at 4°C in sterile PBS containing penicillin and streptomycin, and used within 24 hours. Cords were rinsed in 70% ethanol and sterile PBS and then vein lumens were washed with PBS to remove remaining blood and clots. Cells were released by incubation for 30 minutes at room temperature (RT) in sterile PBS containing calcium, magnesium, and 0.2% collagenase, pH 7.4, with gentle massaging to release cells and then collected in growth media (phenol red–free EBM2 media supplemented with EGM-2 Growth Medium 2 Supplement Mix, penicillin, streptomycin, and amphotericin B). Cells were then centrifuged at 300g for 10 minutes at RT and resuspended in fresh growth media. This suspension was then plated in plastic culture flasks precoated with 0.1% gelatin. Cells were passaged approximately 6-8 days later, upon reaching initial sub-confluence, and then passaged 3 times per week in the absence of antibiotics or frozen in liquid nitrogen until use. Identity and purity of the HUVEC isolations were confirmed each time by more than 99% positive immunostaining with endothelial cell markers (FITC-Ulex europaeus lectin, VE-cadherin) and more than 99.9% negative for α–smooth muscle actin. Cells were assayed between passages 3 and 8. Unless otherwise stated, for all assays, cells were plated at full confluence at a density of 8 × 10^4^ cells/cm^2^ on plates precoated for 30 minutes with 0.1% gelatin and incubated for at least 48 hours prior to the start of experiments.

To stimulate cytokine pathways, HUVECs were treated with 2 μg/ml LPS, 10 ng/ml TNF or 200 ng/mL IL-6 in the presence or absence of 100 ng/mL sIL-6Rα. HUVECs were treated with inhibitors to JAK activity (2 μm ruxolitinib) or p38 (2 μm SB203580) dissolved in DMSO at a maximum concentration of 0.1%, or IKKα/β (5-30 μm BMS345541) dissolved in PBS. A similar amount of DMSO or PBS was added to vehicle control wells.

### Lentiviral Delivery

The lentiviral construct to express SOCS3 with C-terminal Myc and DDK tags was described previously (12). The empty vector used as control was pLenti-C-Myc-DDK-P2A-tGFP. Lentiviral particles were grown in HEK293FT cells by co-transfecting cells with the pLentiC plasmids with pCMV-dR8.2 dvpr and pCMV-VSVG packaging plasmids (18). After 48 hours, medium was concentrated using a 30 kDa cutoff filter, aliquoted, and stored at –80°C until use. Viral titer was determined by the proportion of infected cells with green fluorescence 72 hours post infection.

### Measurement of monolayer permeability

Monolayer permeability was determined in real time by measuring changes in electrical resistance using Electrical Cell-Substrate Impedance Sensor (ECIS, Applied BioPhysics). HUVEC were seeded at confluence onto either an 8W10E or 96W10idf cultureware pre-coated with 0.1% gelatin. Following treatments, the electrical impedance across the monolayer was measured at 1 V, 4000 Hz AC for at least 24 continuous hours and then used to calculate resistance by the manufacturer’s software. Data is presented as a plot of electrical resistance versus time.

### Gene knockdown

Small interference RNA (siRNA) against SOCS3 oligonucleotides were obtained as a set of individual siRNAs from Dharmacon (On target plus SiRNA). Sequences are provided in the supplemental table 3. Cells were transfected with individual siRNA complexed with Lipofectamine siRNA iMAX (Invitrogen) in suspension and seeded at 10^5^ cells/cm^2^. Controls were transfected with on target plus non-targeting control pooled duplexes (Horizon Discovery). Knockdown efficiency was determined by Western blot analysis.

### Gel Electrophoresis and Immunoblotting

Cells grown on 12 well plates were scraped after lysis in 150 μl Laemmli buffer containing the following protease and phosphatase inhibitors: Complete protease inhibitor mixture (Roche Applied Science), PhosSTOP phosphatase inhibitor mixture (Roche Applied Science), 0.1 M NaF, 0.1 mM phenylarsine oxide, 10 mM pyrophosphate and 0.1 mM pervanadate (Sigma-Aldrich). After boiling, a total of 15 μl of cell lysate per lane was loaded on standard SDS-PAGE gels and transferred to nitrocellulose membranes (Bio-Rad). Immunoblots were performed by blocking the membranes with 5% nonfat dry milk or 5% BSA in PBS/Tween and incubating overnight at 4 °C with respective primary antibodies (corresponding dilutions are provided in the supplemental table 2). Secondary HRP-conjugated anti-mouse, anti-rabbit, or anti-goat antibodies were incubated for 1 h at room temperature. Membranes were developed using Clarity or Clarity Max (Bio-Rad) chemiluminescent substrates and a Chemidoc MP Imaging System (BioRad).

### Immunofluorescence Microscopy

Immunofluorescence studies were performed by seeding 8×10^4^ cells/well on 8-well μ-slide chambers (Ibidi) precoated with 0.1% gelatin. At 48 hours post seeding, cells were treated as indicated. Cells were then fixed with 4% paraformaldehyde (Affymetrix) in PBS for 30 min at 4°C, washed twice with PBS, and processed for immunofluorescence at room temperature. Briefly, cells were permeabilized with 0.1% Triton X-100 (Sigma) in PBS (PBS-TX) for 15 min, blocked with 5% bovine serum in PBS-TX. Antibodies were incubated for 2 h at room temperature. After washes in PBS-TX, secondary Alexa Fluor 647-conjugated anti-rabbit and Alexa 594-conjugated anti-goat (Invitrogen) were incubated for 1 h at room temperature in the presence of 4,6-diamidino-2-phenylindole (DAPI, Invitrogen). Images were obtained using a Leica Thunder microscope and analyzed using the manufacturer’s software. Junctional and non-junctional ZO-1 intensity levels were measured in Fiji/ImageJ (19, 20) by creating a binary mask and selection based on the VE-cadherin staining and measuring the intensity of ZO-1 staining in ROIs based on the selection (junctional) and the inversed selection (non-junctional). ZO-1 breaks were counted manually with automated brightness enhancement. Two or more breaks on the same cell-cell border were counted as a single break.

### RT-qPCR

Cells grown on multiwell plates were lysed with Trizol reagent (Thermo Fisher) and total RNA was isolated following manufacturer’s instructions. 400 ng of total RNA was used to prepare cDNA using Primescript RT Master Mix at 42 °C (Clontech) following manufacturer’s instructions. Depending on the target, cDNA was diluted 10-fold in nuclease-free water. Then, 2 μl of cDNA was used per PCR reaction. qPCR was performed in a StepOnePlus (Applied Biosystems) instrument using SYBR green-based iTaq supermix (Bio-Rad) and 2 pmol primers (Thermo Fisher). Fold induction was calculated using the ΔΔCt method using GAPDH as housekeeping gene. Primer sequences utilized are listed in the supplemental table 4.

### Statistics

All statistical analysis and graphs were made in GraphPad Prism version 6. Analysis for RNA and protein expression levels was performed using 1-way ANOVA and Dunnett’s post hoc test or a 2-way ANOVA and Holm-Šídák post hoc tests comparing all samples versus a control (designated in each figure). Two-group comparisons were made with 2-tailed Student’s t test. A 2-tailed P value of less than 0.05 was considered significant. For Western blot experiments, normalized band intensity values (protein/actin) were used. ECIS data were analyzed by 2-way ANOVA of repeated measurements and post hoc analysis comparing main column (treatment) effects.

## Results

LPS-mediated activation of NF-κB leads to increased IL-6 expression (11, 12). Thus, we first tested whether inhibition of signaling downstream of IL-6 receptor prevented LPS effects in endothelial cells. Consistent with prior reports, treatment of HUVEC with LPS leads to a modest and transient increase in monolayer permeability (Figure 1A) despite robust signaling leading to NF-κB and p38 activation (Figure 1B). Notably, LPS treatment led to a small increase in STAT1 and STAT3 phosphorylation that did not achieve statistical significance (Figure 1C), despite increases in IL-6 and SOCS3 mRNA levels (Figure 1D). Notwithstanding, inhibition of STAT phosphorylation by pretreatment with the JAK inhibitor ruxolitinib abrogated the barrier function loss observed after LPS (Figure 1A). This treatment inhibited LPS-induced STAT1/3 phosphorylation (Figure 1C) and the LPS-induced increase in SOCS3 expression, without interfering with the increases in TNF or ICAM1 (Figure 1D), suggesting that ruxolitinib acted upstream of STAT signaling.

**Figure 1:**
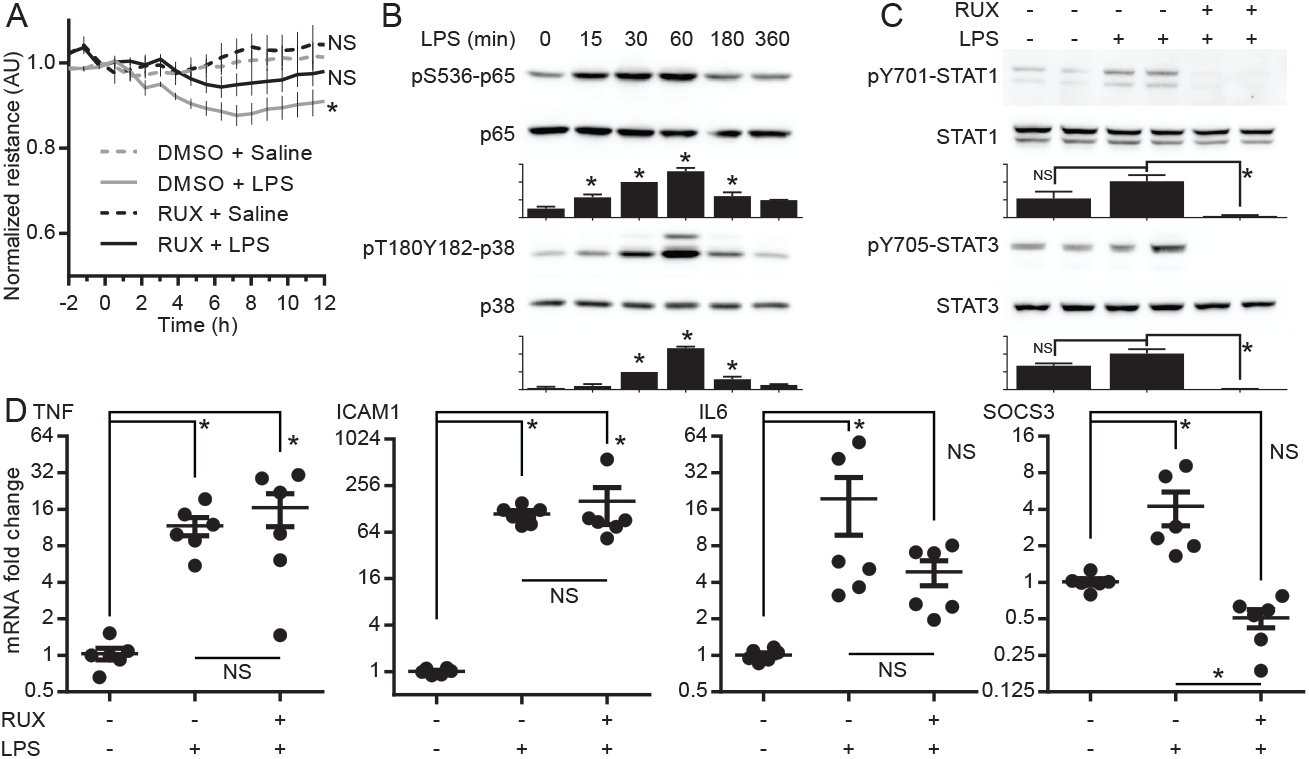
JAK signaling inhibition prevents the mild loss of barrier function induced by endotoxin. *A*: Confluent HUVEC were grown on ECIS arrays. Cells were treated with 2 μM ruxolitinib (RUX) or 0.1% DMSO (as vehicle control). 2 μg/ml LPS was added 30 min after the inhibitor and TEER was measured at 4 kHz every 5 min for at least 24 h. Asterisk, p<0.05 TEER in DMSO+LPS-treated cells vs DMSO+PBS. NS: non-significant differences vs DMSO+PBS. Two-way ANOVA of repeated measurements with Sidak post-hoc tests (n=7-13 from five independent experiments). *B*: Western blots of confluent cells treated with LPS for the indicated times to measure phosphorylated and total levels of p65 (Ser536) or p38 (T180Y182). Ratios of phosphorylated/total proteins were calculated and shown as mean ± SEM and normalized to the 30 min LPS band intensity (n=3 independent experiments). *C*: Confluent cells were treated for 30 minutes with 2μM RUX or 0.1% DMSO, followed by 2 μg/ml LPS or PBS. Cells were lysed 6 hours post LPS treatment to measure phosphorylated and total levels of STAT1 (Y701) and STAT3 (Y705) by Western blot. Ratios of phosphorylated/total proteins were calculated and shown as mean ± SEM (n=8 from 4 independent experiments). *D*: Cells were treated as in C and total RNA was extracted 6 h after LPS treatment. RT-qPCR was performed to measure TNF, ICAM1, IL6 and SOCS3 mRNA expression levels. *GAPDH* was used for normalization and expressed as fold change vs control. Statistics for B-D by one-way ANOVA and Sidak post-hoc tests. Asterisks denote p<0.05 and NS denotes a non-significant change in the comparisons marked in the respective panels.

Endothelial cells express very low levels of IL-6Rα, and addition of recombinant sIL-6Rα is required for full IL-6 signaling (9). We thus hypothesized that lack of this receptor subunit could explain the mild STAT1/3 phosphorylation despite a strong induction of IL-6 expression. To directly test this, we cotreated HUVEC with LPS and sIL-6Rα (herein, LPS+R). This co-treatment led to strong STAT1/3 phosphorylation 3-6 h post-treatment (Figure 2A) and dramatically impaired HUVEC monolayer resistance (Figure 2B). We obtained similar results when adding sIL-6Rα to TNF-treated HUVEC (Figures 2C and 2D). Consistent with low levels of basal IL-6 production by these cells, addition of sIL-6Rα alone did not significantly alter HUVEC monolayer permeability (Figures 2B and 2C). In addition, treatment of HUVEC monolayers with sIL-6Rα did not significantly increase the phosphorylation levels of STAT1 or STAT3 (Figure 2E). An IL-6 amplifier loop maintains high levels of circulating IL-6 through a strong positive feedback loop in severe inflammatory reactions (5). As shown in Figure 2F, LPS+R cells showed a sustained increase in IL-6 mRNA levels, as opposed to the lower and transient increase mediated by LPS alone. Consistent with sustained IL-6 autocrine signaling, LPS+R led to a >60-fold induction that lasted for at least six hours, while LPS alone induced only a modest increase in the levels of SOCS3. Moreover, the expression levels of two markers of an inflamed endothelium, PTGS2 (which codes for COX-2) and F3 (coding for tissue factor, TF), were significantly increased with LPS+R treatments when compared to LPS alone (Figure 2G).

**Figure 2:**
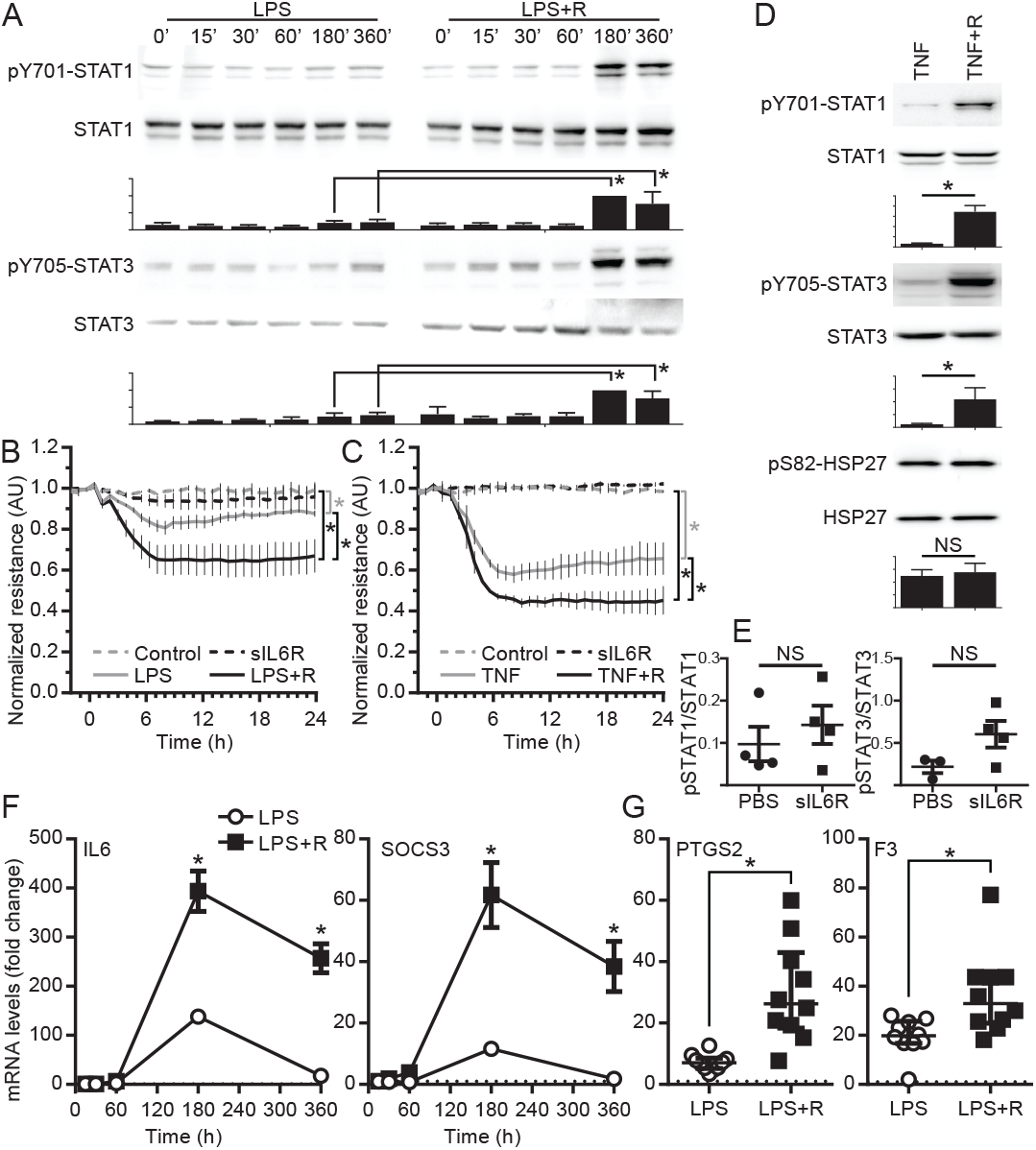
Recombinant sIL6Rα potentiates the response to LPS. *A*: Confluent cells were treated with LPS in the presence (LPS+R) or absence (LPS) of sIL6Rα for indicated times prior to lysis. Phosphorylated and total levels of STAT1 (Y701) and STAT3 (Y705) were measured by Western blot. PBS was used as vehicle control. Ratios of phosphorylated/total proteins were calculated and shown as mean ± SEM and normalized to the 3h LPS+R band intensity. Asterisks denote p<0.05 vs control (no sIL6Rα). Two-way ANOVA with Sidak post-hoc tests (n=4 independent experiments). *B-C*: Confluent HUVEC were grown on ECIS arrays. Cells were treated with either LPS (B) or recombinant TNFα (C) in the presence or absence of sIL6Rα. TEER was measured at 4 kHz every 5 min for at least 24hrs and shown as mean ± SEM (B, n=5 and C, n=3 independent experiments). Two-way ANOVA of repeated measurements with Sidak post-hoc test. Asterisks denote p<0.05 of comparisons marked in the respective panels. *D*: Confluent cells were treated with TNFα or TNFα+sIL6Rα (TNF+R) for different times as indicated prior to lysis. Phosphorylated and total levels of STAT1 (Y701), STAT3 (Y705), or HSP27 (S82) were measured by Western blot. PBS was used as vehicle control. Ratios of phosphorylated/total proteins were calculated and shown as mean ± SEM. Asterisks denote p<0.05 vs control (no sIL6Rα). Student’s T test (n=4 independent experiments). *E*: Graphical representation of the quantification of Western blot analysis to measure phosphorylated and total levels of STAT1 (Y701) and STAT3 (Y705) of cells treated or not with sIL6Rα for six hours. NS: nonsignificant changes. Student’s T test (n=4 independent experiments). *F*: Confluent cells were treated with LPS or LPS+R for different times as indicated prior to total RNA extraction. IL6 and SOCS3 expression levels were measured by RT-qPCR. *GAPDH* was used for normalization. Mean ± SEM fold change expressed vs PBS-treated cells (n=6 from three independent experiments). Two-way ANOVA with Sidak post-hoc test. Asterisks denote p<0.05 LPS+R vs LPS at the corresponding time point. *G*: PTGS2 and F3 expression levels were measured in cells treated for 6 h with LPS or LPS+R (or PBS as vehicle control). Mean ± SEM fold change expressed vs PBS-treated cells (n=10 from seven independent experiments). Asterisks denote p<0.05, Student’s T test.

The loss of barrier function induced by IL-6+R in HUVEC correlates with ZO-1 delocalization, without significantly reducing the levels of ZO-1 expression (9). Notably, we observed a similar level of loss of junctional ZO-1 upon LPS or LPS+R treatments (Figures 3A and 3B). Overall, LPS treatment led to an increase in ZO-1 breaks that was markedly aggravated in LPS+R-treated cells (Figure 3B). In contrast with IL-6+R treatment, LPS led to a transient reduction of total ZO-1 protein levels that did not reach statistical significance and that was not altered by co-treatment with sIL-6Rα (Figure 3C), suggesting that ZO-1 loss from cell-cell junctions, rather than loss of protein expression, is responsible for the IL-6-mediated loss of barrier function initiated by an LPS challenge. Together, the findings shown in these three figures suggest that lack of IL-6Rα receptor is a limiting factor in LPS- and TNF-induced endothelial activation and barrier function loss.

**Figure 3:**
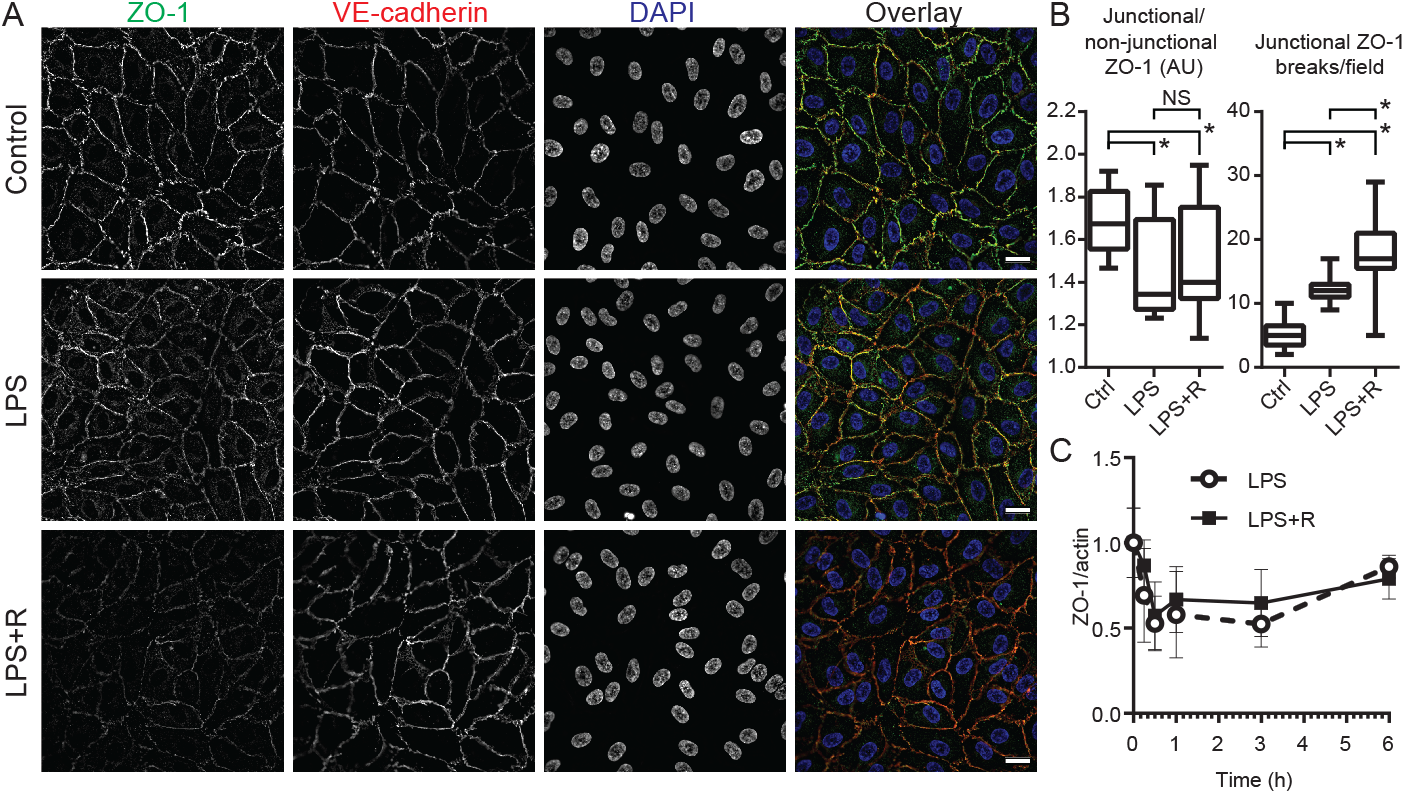
LPS+R treatment leads to a loss of junctional ZO-1. *A*: Confluent HUVEC were treated with vehicle control or LPS in the presence (LPS+R) or absence (LPS) of sIL6Rα for 6hrs. After fixation, cells were stained for VE-cadherin (red) and ZO-1 (green). DAPI (blue) was used to label nuclei. Images are representative fields from three independent experiments. *B*: Quantification of junctional and non-junctional ZO-1 intensity levels as well as the number of junctional ZO-1 breaks per field from cells shown in A (Box and whiskers plot comprising the full data range). Asterisks denote p<0.05. NS: nonsignificant difference. Kruskal-Wallis test with Dunn’s post-hoc comparisons (n=16-25 from three experiments performed in duplicate each, 3-4 random fields per well). C: Levels of ZO-1 protein as determined by Western blot of cells treated with LPS or LPS+R for different times as indicated prior to lysis. Band intensity was normalized to β-actin and shown as mean ± SEM (n=3 independent experiments). Non-significant changes as determined by two-way ANOVA.

We then sought to determine the mechanism underlying the requirement for IL-6Rα. LPS and TNF induce a strong p38 activation, and this pathway has been shown to regulate IL-6 mRNA stability (21), likely affecting any potential autocrine signaling. Thus, we assessed whether altered p38 signaling mediated the autocrine IL-6 signaling effects. We found that sIL-6Rα supplementation to LPS did not affect LPS-induced p38 activation (Figure 4A). Similarly, pharmacologic inhibition of p38 by pretreatment with SB203580 did not alter LPS- or LPS+R-induced barrier function loss (Figure 4B) nor the LPS+R-induced STAT1/3 phosphorylation, despite effectively blunting the phosphorylation of the p38 downstream target HSP27 (Figure 4C). We concluded that p38 activation was not involved in the IL-6 autocrine signaling-induced barrier function loss downstream of LPS in HUVEC.

**Figure 4:**
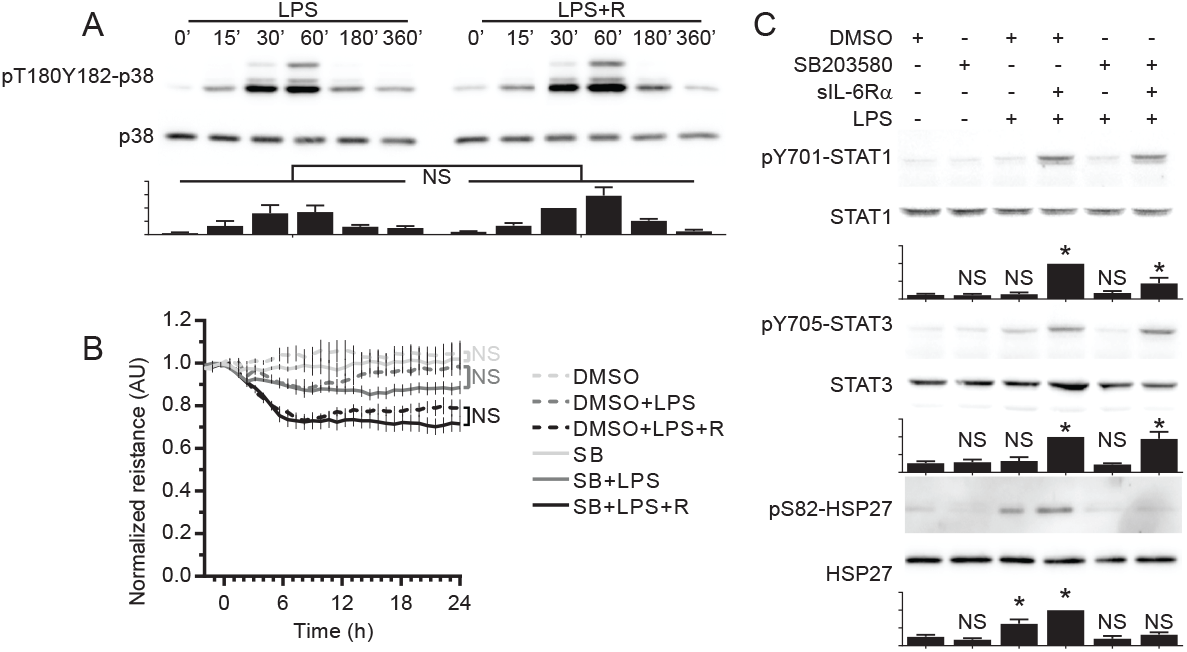
p38 activity is not involved in the barrier loss induced by LPS and sIL6Rα. *A*: Confluent HUVEC were treated with PBS or LPS in the presence (LPS+R) or absence (LPS) of sIL6Rα for the indicated times prior to cell lysis. Phosphorylated and total levels of p38 (T180Y182) were measured by Western blot. PBS was used as vehicle control. Ratios of phosphorylated/total proteins were calculated and shown as mean ± SEM and normalized to the 30 min LPS+R band intensity. NS: non-significant differences in LPS+R vs LPS. Two-way ANOVA (n=4 independent experiments). *B*: Confluent cells were grown on ECIS arrays. Cells were treated for 30 min with 2 μM SB203580 or 0.01% DMSO (as vehicle control). PBS, LPS or LPS + R were then added, and TEER was measured at 4 kHz every 5 min for at least 24 h. Two-way ANOVA of repeated measurements with Sidak post-hoc test (n=3-6 from three independent experiments). NS: nonsignificant differences in post-hoc comparisons marked. *C*: Confluent cells were treated for 30 min with SB203580 or DMSO followed by treatment with PBS, LPS or LPS + R for 6 h and lysis. Phosphorylated and total levels of STAT1 (Y701), STAT3 (Y705) and HSP27 (S82) were measured by Western blot. Ratios of phosphorylated/total proteins were calculated and shown as mean ± SEM and normalized to the LPS+R band intensity. NS: non-significant differences in LPS+R vs LPS. Two-way ANOVA (n=4 independent experiments).

Co-treatment of LPS with sIL-6Rα did not alter LPS-induced levels of p65 phosphorylation, further supporting the idea that IL-6Rα does not affect NF-κB activity (Figure 5A). To confirm this, we treated cells with BMS345541 (22) to inhibit IKK-mediated NF-κB activation downstream of LPS or LPS+R. As expected, this inhibitor blocked LPS-induced IL-6 mRNA expression (Figure 5B), resulting in the loss of the IL-6 amplification, as measured by SOCS3 and PTGS2 expression (Figure 5B) and STAT1 and STAT3 phosphorylation (Figure 5C). As expected, BMS345541 treatment completely blocked the LPS-induced increase in ICAM1 expression levels (Figure 5B).

**Figure 5:**
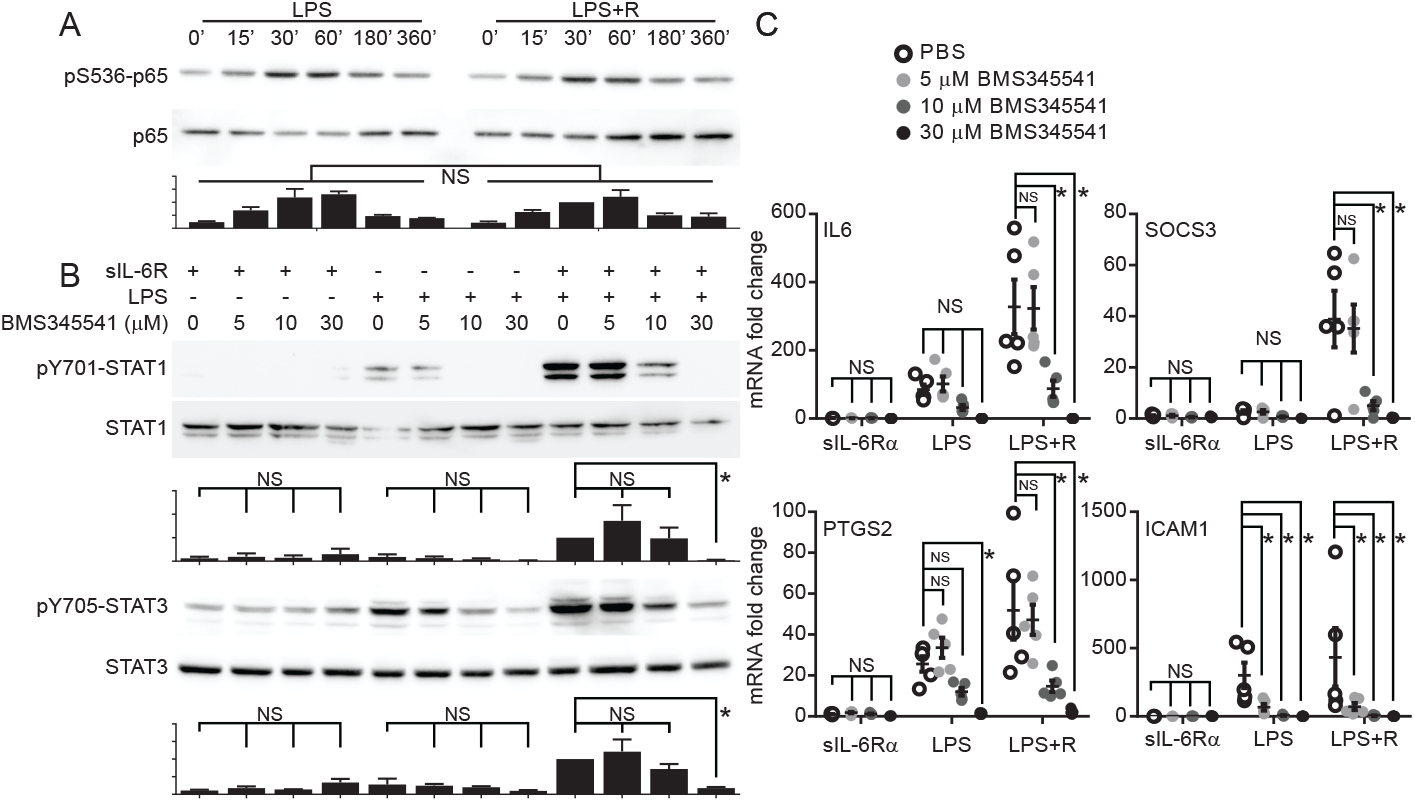
NF-κB activation is required for the initiation of the IL-6 autocrine loop induced by LPS. *A*: Confluent HUVEC were treated with PBS or LPS in the presence (LPS+R) or absence (LPS) of sIL6Rα for the indicated times prior to cell lysis. Phosphorylated and total levels of p65 (S536) were measured by Western blot. PBS was used as vehicle control. Ratios of phosphorylated/total proteins were calculated and shown as mean ± SEM and normalized to the 30 min LPS+R band intensity. NS: non-significant differences in LPS+R vs LPS. Two-way ANOVA (n=4 independent experiments). *B*: Cells were treated with the indicated amounts of BMS345541 for 30 min prior to addition of PBS, LPS, or LPS+R. Cells were lysed 6 h post-treatment and phosphorylated and total levels of STAT1 (Y701) and STAT3 (Y705) were measured by Western blot. Ratios of phosphorylated/total proteins were calculated and shown as mean ± SEM and normalized to the LPS+R band intensity. Two-way ANOVA with Sidak post-hoc test. Asterisks denote p<0.05 and NS denotes a non-significant change of the comparisons marked in the corresponding panels (n=3 independent experiments). *C*: Cells were treated as in B prior to total RNA extraction. IL6, SOCS3, PTGS2 and ICAM1 expression levels were measured by RT-qPCR. *GAPDH* was used for normalization. Mean ± SEM fold change expressed vs PBS-treated cells. Two-way ANOVA with Sidak post-hoc test. Asterisks denote p<0.05 and NS denotes a non-significant change of the comparisons marked in the corresponding panels (n=5 from three independent experiments).

The extent of IL-6 effects on endothelia are mainly limited by a SOCS3-mediated negative feedback loop (5, 10, 12). Thus, we sought to determine if SOCS3 expression in HUVEC regulates the loss of barrier function downstream of LPS. Consistent with a very modest signaling activation of the STAT3 pathway by LPS, SOCS3 knockdown did not significantly modify LPS-induced barrier breakdown (Figure 6A). In contrast, this knockdown further increased the barrier function loss induced by LPS+R co-treatment (Figure 6B), further implicating this pathway in vitro and mirroring our earlier findings in endotoxemic mice (12). We then sought to determine whether inhibition of JAK signaling via SOCS3 overexpression was able to limit the effect of this IL-6 autocrine loop. To test that, we treated cells overexpressing SOCS3 (or control cells transduced with an empty vector) with LPS or LPS+R. As shown in Figure 7A, overexpression of SOCS3 did not inhibit the small barrier function loss induced by LPS treatment, but completely abrogated the increased loss induced by LPS+R, again supporting the notion that simultaneous TLR4 and IL-6 receptor signaling are required for maximal barrier function loss. RT-qPCR data confirmed robust SOCS3 overexpression in all experimental groups (using primers against the SOCS3 CDS, detecting total levels of SOCS3 mRNA derived from both endogenous and exogenous mRNA) and strong pathway inhibition as shown by reduced expression of LPS+R-induced endogenous SOCS3 as measured by primers against the SOCS3 3’ UTR to detect only endogenous mRNA levels (Figure 7B). Similarly, SOCS3 overexpression was able to block the increased barrier function loss induced by TNF+R, although this inhibition was lost at later time points despite robust inhibition of STAT3 transcriptional activity (Figures 7C and 7D). SOCS3 overexpression also inhibited sIL-6Rα-induced increases in LPS-induced induction of PTGS2 and F3 expression, but not the NF-κB target ICAM-1 (Figure 7E). Lastly, we found that JAK inhibition rescued the barrier function loss induced by LPS+R even when added 8h post-LPS+R treatment (Figure 7F), demonstrating that continued JAK signaling is required downstream of LPS+R and suggesting that an IL-6 amplifier loop, similar to the loop described in vivo (5), is active in cultured HUVEC and responsible for the sustained barrier function loss.

**Figure 6:**
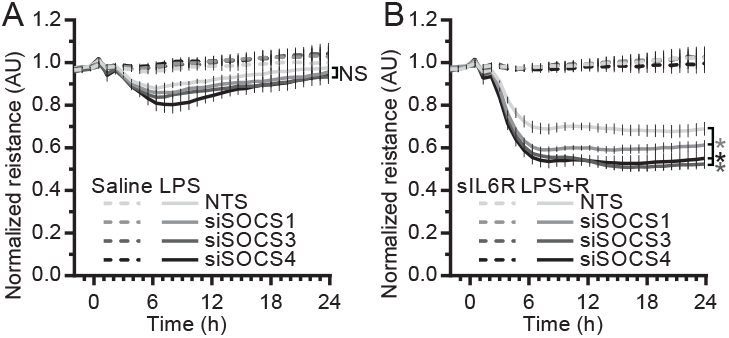
SOCS3 knockdown further impairs barrier function of HUVEC treated with LPS+R. HUVEC were transfected with 50 nM of one of three independent SOCS3 siRNA sequences (or a non-targeting sequence, NTS) and then plated on ECIS arrays. Cells were allowed to grow to confluency and achieve a steady state resistance. Then, cells were treated with either PBS or LPS (A), or sIL6Rα or LPS+R (B). TEER was measured at 4 kHz every 5 min for at least 24 h and shown as mean ± SEM normalized to the measurement immediately prior to treatment (n=6 from three independent experiments performed in duplicate). Two-way ANOVA of repeated measurements and Sidak post-hoc tests. NS denotes nonsignificant changes and asterisks denote p<0.05 of the respective siRNA vs NTS in cells treated with LPS (A) or LPS+R (B).

**Figure 7:**
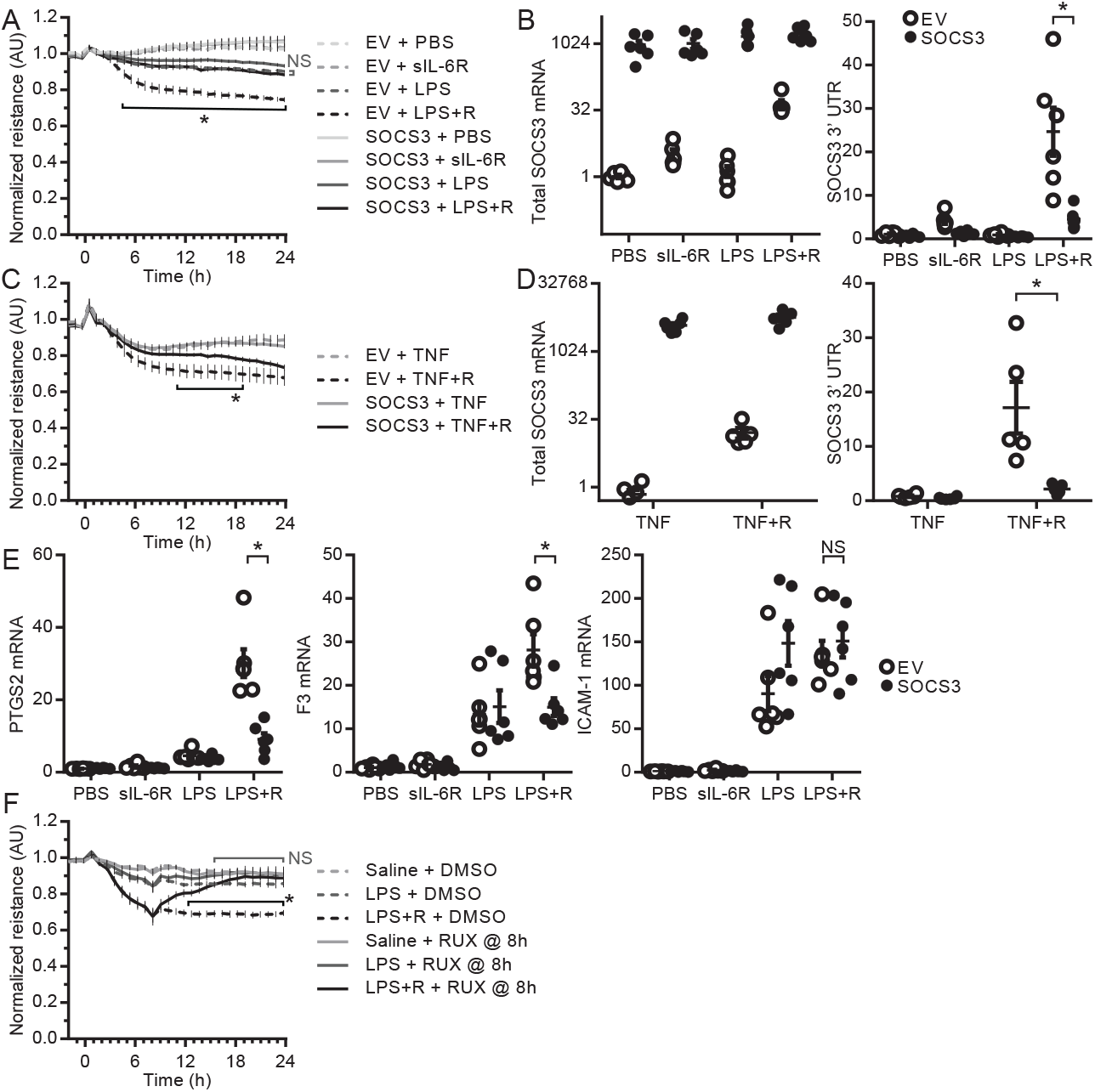
SOCS3 overexpression prevents the increased barrier function loss induced by LPS+R and JAK inhibition restores barrier function. *A*: HUVEC transduced with lentivirus to overexpress SOCS3 or with an empty vector (EV) control were seeded onto ECIS arrays and allowed to grow to confluency. Cells were then treated with either PBS, sIL6Rα, LPS or LPS+R. TEER was measured at 4 kHz every 5 min for at least 24 h and shown as mean ± SEM normalized to the measurement immediately prior to treatment (n=11 from four independent experiments). *B*: Total RNA from cells treated as in A was extracted 6 h post-treatment. RT-qPCR was performed with primers against the SOCS3 CDS (detecting total levels of SOCS3 mRNA derived from both endogenous and exogenous mRNA) and against the SOCS3 3’ UTR (to detect only endogenous mRNA levels). *GAPDH* was used for normalization. Fold change vs EV+PBS shown as individual values and mean ± SEM (n=6 from three independent experiments). C: HUVEC transduced as in A were treated with TNF in the presence (TNF+R) or absence (TNF) of sIL6Rα. TEER was measured at 4 kHz every 5 min for at least 24 h and shown as mean ± SEM normalized to the measurement immediately prior to treatment (n=9-10 from four independent experiments). *D*: Total RNA from cells treated as in C was extracted 6 h post-treatment. RT-qPCR was performed to detect total and endogenous SOCS3 as in B. Fold change vs EV+PBS shown as individual values and mean ± SEM (n=6 from three independent experiments). *E*: PTGS2, F3 and ICAM1 expression levels were measured by RT-qPCR from cells treated as in B. *GAPDH* was used for normalization. Fold change vs EV+PBS shown as individual values and mean ± SEM (n=6 from three independent experiments). *F*: Confluent HUVEC were grown on ECIS arrays. Cells were treated with PBS, LPS or LPS+R for 8 h to allow for stable maximal loss of barrier function, followed by a treatment with 2 μM ruxolitinib (RUX) or 0.01% DMSO (as vehicle control). TEER was measured at 4 kHz every 5 min for at least 24 h and shown as mean ± SEM normalized to the measurement immediately prior to treatment (n=7-8 from three independent experiments). A, C, F: Two-way ANOVA of repeated measurements with Sidak post-hoc test. Asterisks denote p<0.05 and NS denote a non-significant difference of the comparisons made as marked in the respective panels. A horizontal bracket denotes statistical significance (or lack of) during the marked period. B, D, E: Two-way ANOVA with Sidak post-hoc test. Asterisks denote p<0.05 and NS denote a nonsignificant difference of the comparisons made as marked in the respective panels.

## Discussion

Acute inflammation leads to a large increase in the levels of circulating cytokines, resulting in several endothelial responses, including increases in leukocyte binding, thrombosis, and vascular permeability (1–4). The effects of the signaling induced by individual cytokines is well understood. However, little is known about the endothelium response when exposed to multiple simultaneous signals. Furthermore, soluble receptors may act as co-agonists or traps for the circulating cytokines, adding an additional level of complexity to cytokine signaling in the context of acute inflammation. The circulating levels of interleukin 6 (IL-6) correlate with systemic disease severity (5, 23) and are highly predictive of mortality (24, 25). In fact, an ‘IL-6 amplifier’ mechanism that consists in a positive feedback regulation of this cytokine has been linked to worse prognosis (5). However, the clinical failure of anti-inflammatory drugs and monoclonal antibodies to treat septic patients over the last two decades (26) demonstrates that blocking circulating proinflammatory signals is not enough to improve the condition of these patients. After decades, even the utility of corticosteroids is being debated (27). Direct inhibition of the mechanisms within the endothelium responsible for vascular dysfunction in response to proinflammatory signaling provide a potential avenue to prevent or even reverse shock-induced endotheliopathy, thus improving the prognosis of severely ill patients without reducing the immune response necessary for pathogen clearance.

Here, we present evidence that in the presence of sIL-6Rα, NF-κB activation in cultured endothelial cells promotes an increase in IL-6 expression and autocrine signaling that synergistically leads to the loss of endothelial barrier function and to the expression of pro-inflammatory gene expression. Inhibition of JAK signaling downstream of IL-6R, either by the pharmacological kinase inhibitor ruxolitinib or overexpression of the negative regulator SOCS3, blunts this autocrine signaling, without affecting the direct TLR-4 or TNFR effects. Strikingly, ruxolitinib treatment was able to rescue barrier function in HUVEC even 8 h after LPS+R treatment, demonstrating that continuous JAK activity is required to maintain barrier function loss. This finding is consistent with our prior observations in response to direct IL-6 receptor activation. Together, the data shown here provide direct proof of a critical role for the IL-6/JAK/STAT3/SOCS3 signaling axis in the regulation of endothelial function during acute inflammation and offer mechanistic insight for our prior observation that deletion of SOCS3 expression specifically in the endothelium led to severe endotheliopathy, organ dysfunction and lethality in response to a challenge with LPS that is not lethal in control mice (12). We show that LPS-induced NF-κB activation is not sufficient to induce the full extent of endothelial barrier function loss, and requires IL-6 signaling. SOCS3, as a negative regulator of this pathway, limits this response.

Endothelial cells readily respond to IL-6 stimulation through a trans-signaling mechanism that requires a rise in the levels of soluble forms of IL-6Rα (9, 12). Multiple inflammation-driven mediators secreted by, or expressed at, the endothelial luminal surface, including IL-6 itself, cause systemic vascular leak, coagulopathy, leukocyte adhesion and vasodilation, contributing to organ failure (2–4, 28–31). We recently showed that loss of the IL-6 pathway inhibitor SOCS3 leads to increased severity and acute mortality after an endotoxin challenge that is not lethal in control mice. This response was associated with intravascular leukocyte accumulation, thrombocytopenia, and systemic vascular leak, as well as high levels of circulating IL-6, but not TNF. Gene expression assays in these mice as well as in cultured HUVEC led us to propose that loss of SOCS3 leads to an overactivation of IL-6 signaling that dramatically alters the endothelial transcriptional profile to promote an increased inflammatory response, including a potential autocrine IL-6 feedback loop (12). Here, we devised several experiments to test this hypothesis. For that, we took advantage of the fact that, as most other endothelial cells, HUVEC express one of the IL-6 receptor subunits (gp130), but not the other (IL-6Rα, or gp80) (8, 9). Accordingly, in the absence of recombinant IL-6Rα, the response to IL-6 by HUVEC is barely detectable. This system allowed us to directly test if endogenous levels of IL-6, as induced by NF-κB activation, had a significant effect on endothelial dysfunction. We show that addition of sIL-6Rα to LPS (LPS+R) dramatically increased the response to LPS. We observed a similar synergistic effect when cells were exposed to TNF in the presence of IL-6Rα (TNF+R). Inhibition of JAK activity by ruxolitinib completely abrogated this effect. LPS+R promoted a similar loss of junctional ZO-1 as observed by direct IL-6 receptor activation, suggesting that a common mechanism underlies these responses. These findings show that activation of JAK/STAT signaling is a crucial factor to promote endothelial dysfunction in response to endotoxin and can explain the observation that LPS promotes a strong endothelial dysfunction in animal models but little barrier function loss in cultured endothelial cells. We then went on to show that SOCS3 is a critical regulator of this response. SOCS3 knockdown further increased barrier function loss in HUVEC treated with LPS+R, but not in cells treated with LPS alone. Conversely, HUVEC overexpressing SOCS3 responded to LPS+R by inducing the same level of barrier function loss and gene expression as LPS alone. Together, those findings show that inhibition of JAK activity downstream of IL-6 limits the synergistic effect of STAT and NF-kB activation and provides a mechanism to explain the acute vascular dysfunction and lethality in endotoxemic mice lacking endothelial SOCS3.

Rarely does a patient have the possibility of blocking an inflammatory response prior to its start. Thus, it is critical to find therapeutic strategies that are able to rescue the vascular dysfunction once inflammation occurs. Here, we show that constant JAK activity is required for LPS+R-induced barrier function loss, since its inhibition by ruxolitinib 8 h after treatment, when cells already reached a maximal permeability increase, completely restored barrier function loss. Identification of the mechanisms downstream of JAK activity leading to these changes may result in a successful therapeutic strategy to treat vascular dysfunction in response to proinflammatory cytokines even in those cases in which direct cytokine blockade is not advisable.

## Author contributions

N.M. and A.P.A. conceived and designed research; N.M., R.B.R, D.C., L.T, and A.P.A. performed experiments; N.M., R.B.R, and A.P.A. analyzed data; N.M., R.B.R., D.C., and A.P.A. interpreted results of experiments; N.M., R.B.R., and A.P.A prepared figures; N.M. and A.P.A. drafted manuscript; N.M. and A.P.A. edited and revised manuscript with input from all authors; N.M., R.B.R, D.C., L.T., and A.P.A. approved final version of manuscript.

## Acknowledgements

This project was supported by the National Institute of General Medical Sciences Grant R01GM124133 to A.P.A.

---

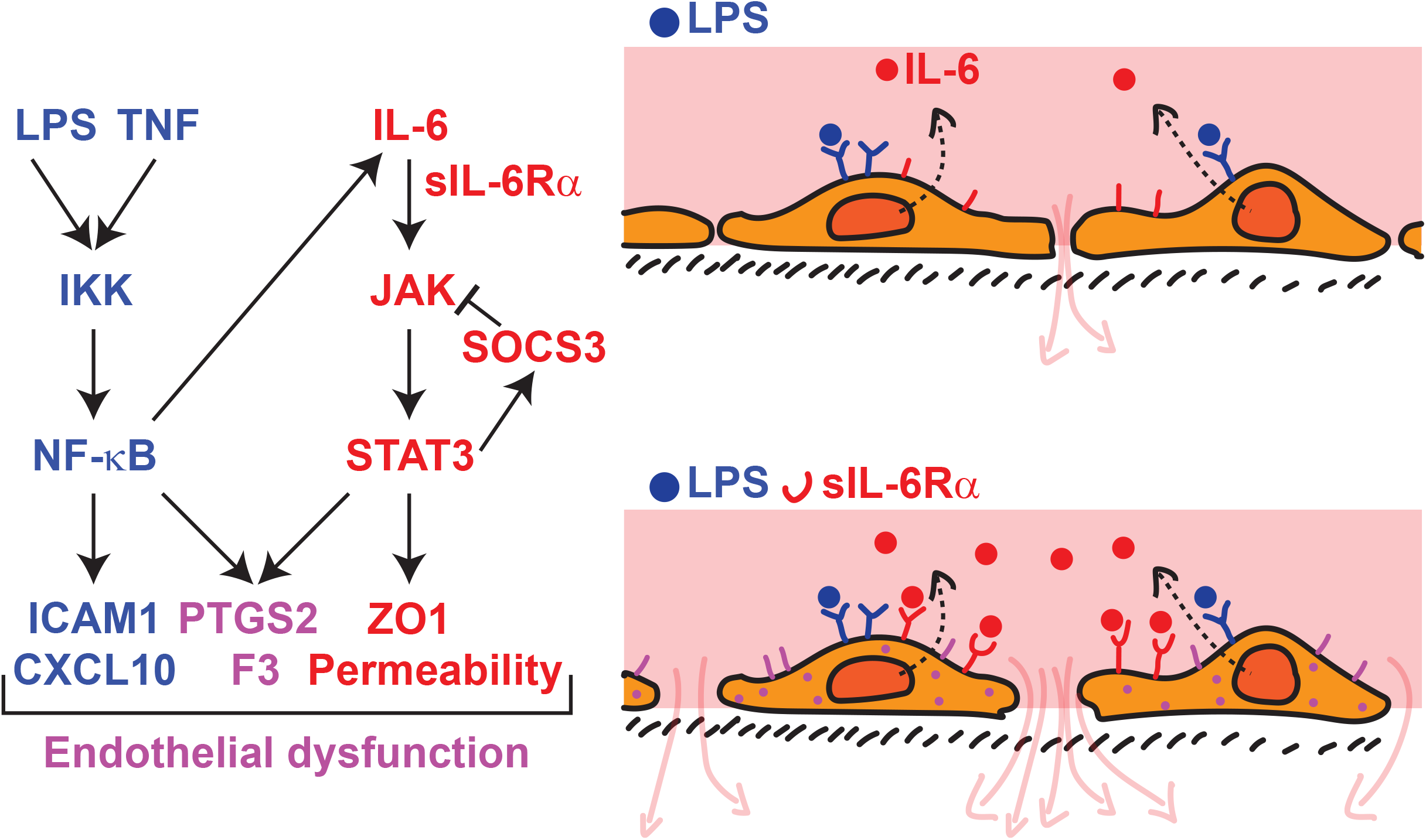

