## Supplemental Tables 1-4 for "SOCS3 limits endotoxin-induced endothelial dysfunction by blocking a required autocrine interleukin 6 signal in human umbilical vein endothelial cells"

| Supplemental Table 1 | | | |
|---|---|---|---|
| **Reagent or disposable** | **Vendor** | **Catalog Number** | **Concentration or dilution** |
| pCMV-dR8.2 dvpr | Addgene | 8455 | N/A |
| pCMV-VSV-G | Addgene | 8454 | N/A |
| 4% PFA | Affymetrix | | N/A |
| Tween 20 | AMRESCO | 0777-1L | 0.1% |
| 8-well electrode array | Applied Biophysics | 8W10E PET | N/A |
| 96-well electrode array | Applied Biophysics | 96W10idf PET | N/A |
| Transblot Turbo RTA Mini Nitrocellulose Transfer kit | Bio-Rad | 1704270 | N/A |
| Clarity Western ECL Substrate | Bio-Rad | 1705061 | As per instructions |
| Clarity Max Western ECL Substrate | Bio-Rad | 1705062 | As per instructions |
| iTaq Universal SYBR Green Supermix | Bio-Rad | 1725125 | 1X |
| SB203580 | Cayman Chemicals | 13067 | 2 µM |
| BMS345541 | Cayman Chemicals | 16667 | 5-30 µM |
| Penicillin-Streptomycin Solution, 100x | Corning | 30-002-CI-PK | 1X |
| Trypsin-EDTA Solution 1X | Corning | 25-052-CI | 1X |
| ON-TARGETplus set of 4 SOCS3 siRNA | Horizon Discovery | LQ-004299-00-0005 | 50 nM (35 pmol siRNA/4 $cm^2$ well) |
| ON-TARGETplus non-targeting control pool | Horizon Discovery | D-001810-10-20 | 50 nM (35 pmol siRNA/4 $cm^2$ well) |
| IBIDI 8 well uslide | IBIDI | 80826-90 | N/A |
| lipofectamine RNAiMAX transfection reagent | Invitrogen | 13778150 | 6 pmol siRNA/µl lipid (6 µl lipid/4 $cm^2$ well) |
| TRIzol reagent | Invitrogen | 15596018 | 1X |
| gelatin | Millipore Sigma | ES-006-B | 0.1% |
| Antibiotic Antimycotic Solution (100×), Stabilized | Millipore Sigma | A5955-20ML | 1X |
| FITC-Ulex europaeus lectin | Millipore Sigma | L9006 | 1:200 |
| Sodium fluoride | Millipore Sigma | S-1504 | 100 mM |
| phenyl arsine oxide | Millipore Sigma | P-3075 | 100 µM |
| Sodium pyrophosphate decahydrate | Millipore Sigma | 221368 | 10 mM |
| Sodium orthovanadate | Millipore Sigma | S6508 | 100 µM |
| Lipopolysaccharides from Escherichia coli O111:B4 (LPS) | Millipore Sigma | L4391 | 2 µg/ml |
| Triton X-100 | Millipore Sigma | T8787 | 0.1% |
| DMSO | Millipore Sigma | 472301 | <0.1% |
| Fetal Bovine Serum | Millipore Sigma | 12306C | 0.05 |

| | | | |
|---|---|---|---|
| pLenti-C-Myc-DDK-P2A-tGFP Lentiviral Gene Expression Vector | Origene | PS100088 | N/A |
| phenol red-free EBM 2 media | PromoCell | C-22216 | N/A |
| EGM-2 Growth Medium 2 Supplement Mix | PromoCell | C-39216 | As per instructions |
| Recombinant human IL-6 | R&D Systems | 206-IL-200/CF | 200 ng/mL |
| Recombinant human sIL-6Rα | R&D Systems | 227-SR-025/CF | 100 ng/mL |
| Recombinant Human TNF-alpha | R&D Systems | 210-TA-020 | 10ng/ml |
| complete protease inhibitor mixture | Roche Applied Science | 11697498001 | 1X |
| PhosSTOP phosphatase inhibitor mixture | Roche Applied Science | 04906837001 | 1X |
| Bovine Serum Albumin | Rockland Immunochemical | BSA-1000 | 3-5% |
| Vivacell 100, 30,000 MWCO PES, 10pc | Sartorius | VC1022 | N/A |
| Ruxolitinib | Selleck Chem | S1378 | 2 μM |
| PrimeScript RT Master Mix | Takara Bio | RR036B | 1X |
| Opti-MEM | Thermo Fisher Scientific | 31985070 | 1X |
| 293FT cell line | Thermo Fisher Scientific | R700-07 | N/A |
| DAPI (4',6-Diamidino-2-Phenylindole, Dilactate) | Thermo Fisher Scientific | D3571 | 1 ug/ml |
| Type 1 Collagenase | Worthington Biochemical | LS004196 | 0.2% |

| Supplemental Table 2 | | | | | | | |
|---|---|---|---|---|---|---|---|
| **Target** | **Species** | **Clone or isotype** | **Vendor** | **Catalog No** | **RRID** | **Concentration or dilution** | **Blocking solution** |
| β-actin | mouse | clone AC-15 | Millipore Sigma | A5441 | AB_476744 | 1:10000 | 5%BSA, 0.1% Tween in PBS |
| HSP 27 | mouse | monoclonal | Cell Signaling Technology | 2402 | AB_331761 | 1:1000 | 3% milk in PBS + 0.1% Tween |
| pS82 HSP27 | rabbit | polyclonal | Cell Signaling Technology | 2401 | AB_331644 | 1:1000 | 3% milk in PBS + 0.1% Tween |
| STAT3 | rabbit | clone D1A5 | Cell Signaling Technology | 8768 | AB_2722529 | 1:5000 | 5%BSA, 0.1% Tween in PBS |
| pY705-STAT3 | rabbit | clone D3A7 | Cell Signaling Technology | 9145 | AB_2491009 | 1:5000 | 5%BSA, 0.1% Tween in PBS |
| SOCS3 | rabbit | polyclonal IgG | Abcam | AB16030 | AB_443287 | 1:2000 | Block for 1 hour- 5% BSA,0.1% Tween in PBS. Primary Ab in PBS ONLY overnight at 4C |
| STAT1 | rabbit | polyclonal | Cell Signaling Technology | 9172 | AB_2198300 | 1:1000 | 5%BSA, 0.1% Tween in PBS |
| pY701-STAT1 | rabbit | clone D4A7 | Cell Signaling Technology | 7649 | AB_10950970 | 1:1000 | 5%BSA, 0.1% Tween in PBS |
| p38 | rabbit | polyclonal | Santa Cruz | sc-535 | AB_632138 | 1:500 | 3% milk in PBS + 0.1% Tween |
| pT180Y182-p38 | rabbit | clone D3F9 | Cell Signaling Technology | 4511 | AB_2139682 | 1:1000 | 5%BSA, 0.1% Tween in PBS |
| p65 | rabbit | clone D14E12 | Cell Signaling Technology | 8242 | AB_10859369 | 1:2000 | 5%BSA, 0.1% Tween in PBS |
| pS536-p65 | rabbit | clone 93H1 | Cell Signaling Technology | 3033 | AB_331284 | 1:2000 | 5%BSA, 0.1% Tween in PBS |
| VE-cadherin | goat | polyclonal | R&D Systems | AF938 | AB_355726 | 1:100 | 5% FBS in PBS |
| ZO-1 (for IF) | rabbit | polyclonal | Thermo Fisher Scientific | 40-2200 | AB_2533456 | 1:200 | 5% FBS in PBS |
| ZO-1 (for WB) | rabbit | clone D6L1E | Cell Signaling Technology | 13663 | AB_2798287 | 1:500 | 5%BSA, 0.1% Tween in PBS |
| Peroxidase AffiniPure Goat | mouse | polyclonal | Jackson Immunoresearch | 115-035-062 | AB_2338504 | 1:5000 | Same as primary Ab blocker |

| | | | | | | | |
|---|---|---|---|---|---|---|---|
| Anti-Mouse IgG (H+L) | | | | | | | |
| Peroxidase AffiniPure Goat Anti-Rabbit IgG (H+L | rabbit | polyclonal | Jackson Immunoresearch | 111-035-003 | AB_2313567 | 1:5000 | Same as primary Ab blocker |
| Peroxidase AffiniPure Bovine Anti-Goat IgG (H+L) | goat | polyclonal | Jackson Immunoresearch | 805-035-180 | AB_2340874 | 1:5000 | Same as primary Ab blocker |
| Donkey anti-Rabbit IgG (H+L) Highly Cross-Adsorbed Secondary Antibody, Alexa Fluor Plus 647 | donkey | polyclonal | Thermo Fisher Scientific | A32795 | AB_2762835 | 1:500 | 5% FBS in PBS |
| Donkey anti-Goat IgG (H+L) Cross-Adsorbed Secondary Antibody, Alexa Fluor 594 | donkey | polyclonal | Thermo Fisher Scientific | A11058 | AB_2534105 | 1:500 | 5% FBS in PBS |

| Supplemental Table 3 | | | |
| --- | --- | --- | --- |
| **siRNA label** | **Sequence** | **Vendor** | **Catalog No** |
| SOCS3 siRNA #1 | CAGCAUCUCUGUCGGAAGA | Horizon Discovery | J-004299-09 |
| SOCS3 siRNA #3 | GCGAUGGAAUUACCUGGAA | Horizon Discovery | J-004299-11 |
| SOCS3 siRNA #4 | ACAGGGCAGUUGUGUGUUG | Horizon Discovery | J-004299-12 |

**Supplemental Table 4**

| **Gene Name** | **Forward Sequence** | **Reverse Sequence** |
|---|---|---|
| GAPDH | GTCTCCTCTGACTTCAACAGCG | ACCACCCTGTTGCTGTAGCCAA |
| TNF | CTCTTCTGCCTGCTGCACTTTG | ATGGGCTACAGGCTTGTCACTC |
| ICAM1 | AGCGGCTGACGTGTGCAGTAAT | TCTGAGACCTCTGGCTTCGTCA |
| IL6 | TACCACTTCACAAGTCGGAGGC | CTGCAAGTGCATCATCGTTGTTC |
| SOCS3 (CDS) | CATCTCTGTCGGAAGACCGTCA | GCATCGTACTGGTCCAGGAACT |
| SOCS3 (3’UTR) | ACTCTGTGCCTCCTGACTAT | AGGCTGAGTATGTGGCTTTC |
| F3 | CAGACTTCACACCTTACCTGGAG | GTTGTTCCTTCTGACTAAAGTCCG |
| PTGS2 | CGGTGAAACTCTGGCTAGACAG | GCAAACCGTAGATGCTCAGGGA |
